# Apicoplast-derived isoprenoids are essential for biosynthesis of GPI protein anchors, and consequently for egress and invasion in *Plasmodium falciparum*

**DOI:** 10.1101/2024.02.14.580402

**Authors:** Michaela S. Bulloch, Long K. Huynh, Kit Kennedy, Julie E. Ralton, Malcolm J. McConville, Stuart Ralph

## Abstract

Glycophosphatidylinositol (GPI) anchors are the predominant glycoconjugate in *Plasmodium* parasites, enabling modified proteins to associate with biological membranes. GPI biosynthesis commences with donation of a mannose residue held by dolichol-phosphate at the endoplasmic reticulum membrane. In *Plasmodium* dolichols are derived from isoprenoid precursors synthesised in the *Plasmodium* apicoplast, a relict plastid organelle of prokaryotic origin. We found that treatment of *Plasmodium* parasites with apicoplast inhibitors decreases the abundance of isoprenoid and GPI intermediates resulting in GPI-anchored proteins becoming untethered from their normal membrane association. Even when other isoprenoids were chemically rescued, GPI depletion led to an arrest in schizont stage parasites, which had defects segmentation and egress. In those daughter parasites (merozoites) that did form, proteins that would normally be GPI-anchored were mislocalised, and when these merozoites were artificially released they were able to attach to but not invade new red blood cells. Our data provides further evidence for the importance of GPI biosynthesis during the asexual cycle of *P. falciparum*, and indicates that GPI biosynthesis, and by extension egress and invasion, is dependent on isoprenoids synthesised in the apicoplast.

**Author summary:** The plastid apicoplast organelle of the malaria parasite *Plasmodium falciparum* has long been recognised as a drug target, however the downstream metabolic pathways have not been fully elucidated. In this study we inhibited apicoplast function in blood-stage *P. falciparum* and following the depletion of essential apicoplast-derived isoprenoids, we observed that these parasites exhaust their supplies of the polyisoprenoid alcohol dolichol. Dolichols form important components of biological membranes and are also required for the synthesis of the major parasite glycoconjugate, glycophosphatidylinositol (GPI) anchors. Concurrent with a reduction in dolichol levels, proteins normally conjugated to GPIs became mislocalised. Severe parasite impairments followed with incomplete membrane segmentation of their daughter merozoites, which could subsequently neither egress nor reinvade host red blood cells. Our data implicates dolichol as an essential parasite metabolite, dependent on normal apicoplast function, and reveals novel roles for GPI anchored proteins. The widespread phenotype following disrupted dolichol synthesis supports aspects of GPI biosynthesis as potential future drug targets.

## Introduction

The most severe human malaria is caused by infection with *Plasmodium falciparum*, a unicellular protist parasite that is spread through the bite of an infected *Anopheles* mosquito. Like most other apicomplexans, *Plasmodium* has a vestigial non-photosynthetic plastid known as the apicoplast. The prokaryotic origins of the apicoplast present differences from the vertebrate host that are attractive drug targets (1). This self-contained organelle retains its own reduced genome, protein synthesis and import pathways. During the intraerythrocytic development cycle (IDC), the apicoplast houses type II fatty acid synthesis and the synthesis of haem and iron-sulphur clusters (2-4). However, the only essential apicoplast-derived metabolites synthesised during the asexual cycle are coenzyme A (whose synthesis can continue in the absence of an intact apicoplast) and isopentenyl pyrophosphate (IPP). IPP is synthesised via a 1-deoxy-D-xyulose 5-phosphate (DOXP) pathway and is the precursor for all other isoprenoid metabolites (4-7). Apicoplast null parasites can survive indefinitely in culture during the IDC if supplied with sufficient exogenous IPP.

Direct inhibition of the DOXP pathway with inhibitors such as fosmidomycin (which targets 1-deoxy-D-xyulose 5-phosphate reductoisomerase) is rapidly lethal, killing parasites within the same IDC in which it is administered (8). Conversely, inhibitors that target plastid housekeeping functions (such as the protein translation inhibitors indolmycin and clindamycin) have a delayed mechanism of indirectly depleting IPP (9-12). Such housekeeping inhibitors cause parasites in the following cycle to inherit a dysfunctional apicoplast with depleted isoprenoid reservoirs. These parasites die from isoprenoid starvation by the second cycle trophozoite stage in a phenomenon known as delayed death (12-14).

Roles for IPP have been conjectured based on analogy to better characterised eukaryotes, but our knowledge of the cellular fate of apicoplast-synthesised IPP is limited. While some IPP serves roles within the apicoplast, at least some IPP is exported into the cytoplasm where it is condensed to generate 15- and 20-carbon farnesyl and geranylgeranyl polyisoprenoid chains (15, 16). These isoprenoids are post-translationally linked to a number of membrane associated proteins, including Rab GTPases which are required for the correct formation of the parasite digestive vacuole, inner membrane complex (IMC) and cytostomal invaginations (15, 17-20). The mislocalisation of these prenylated proteins is likely to be the proximate cause of delayed death of asexual blood stages following inhibition of IPP synthesis. Protein prenylation can be selectively restored by supplementation with the alcohol derivative geranylgeraniol (GGOH), though this only offers a temporary reprieve for the parasite that we refer to here as Partially Rescued Delayed Death (PRDD) (14). These PRDD parasites only mature to second cycle schizonts before stalling and dying (**Fig 1A**). As continuous parasite viability is not restored by GGOH, apicoplast IPP must also be essential for a prenylation-independent process or processes.

**Fig 1.**
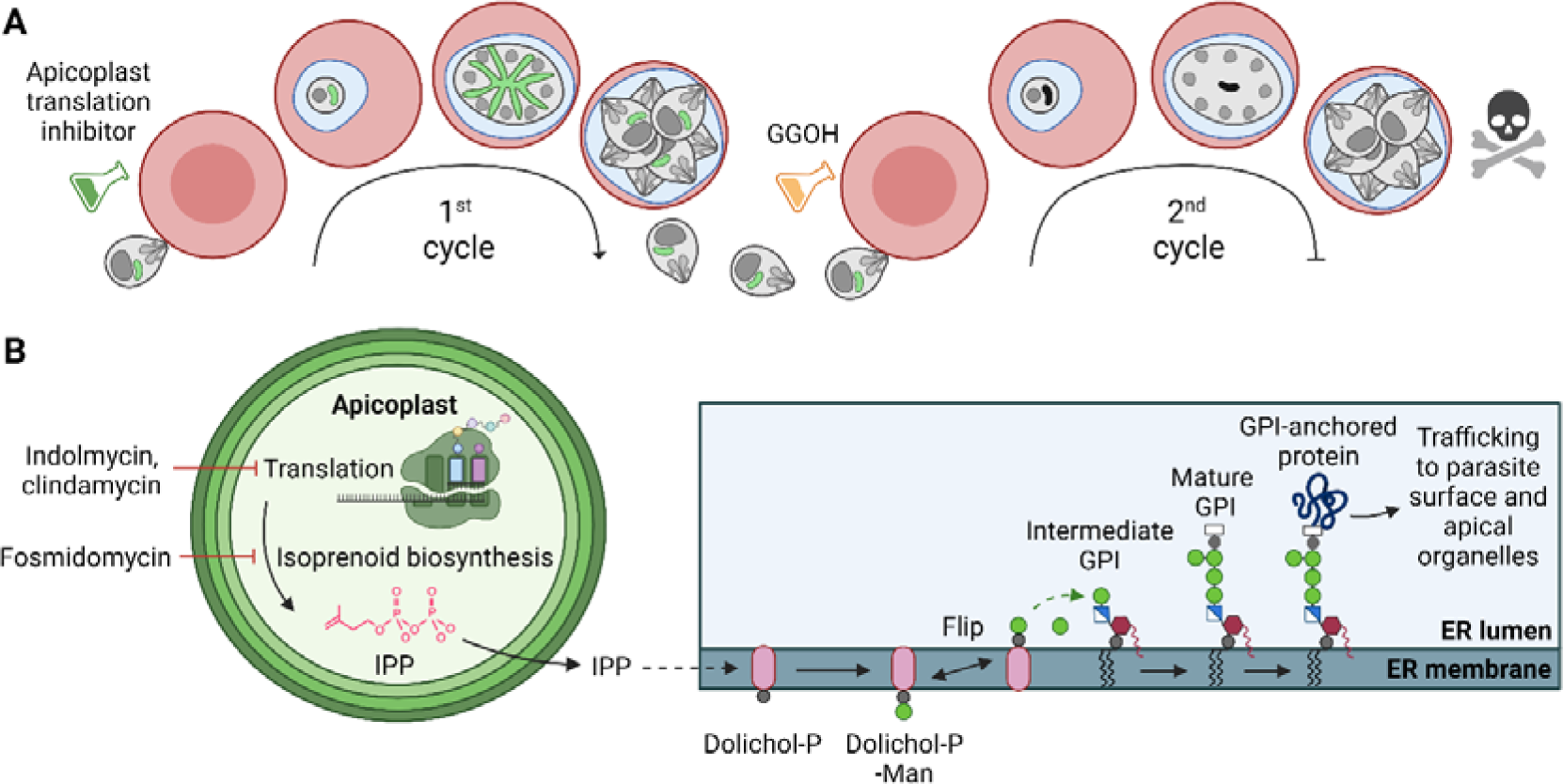
Apicoplast-targeting drugs are lethal to *Plasmodium* parasites in the asexual blood stage due to direct or indirect inhibition of IPP production. **A)** Delayed death is a consequence of inhibition of plastid housekeeping functions in *P. falciparum*. Daughter parasites inherit a dysfunctional apicoplast and so are unable to synthesise the essential metabolite isopentenyl pyrophosphate (IPP), resulting in parasite death only in the cycle following treatment. Supplementation with the prenyl precursor GGOH temporarily rescues delayed death, allowing parasites to progress to the end of the second cycle before succumbing to IPP starvation. **B)** IPP is synthesised in the apicoplast and exported into the cytosol. Multiple IPP units are condensed together to form polyisoprenoids chains which are used for dolichol synthesis in the ER. Dolichol-phosphate-mannose (Dolichol-P-Man) is synthesised at the cytosolic face of the ER, then flipped into the ER lumen, where it donates essential mannose residues to build GPI anchors which are used to tether proteins with glycophosphatidylinositol (GPI) attachments motifs to membranes.

Apicoplast IPP is also expected to be used to synthesise dolichols, very long polyisoprenoid alcohols that are synthesised in the endoplasmic reticulum (ER) via the reduction of polyprenols (**Fig 1B**) (21-23). Dolichol can be phosphorylated to form dolichol-phosphate (dolichol-P) and subsequently dolichol-phospho-mannose (DPM), which acts as a sugar donor for a number of ER mannosyltransferases involved in glycophosphatidylinositol (GPI) biosynthesis and protein tryptophan C-mannosylation (24, 25). The C-mannosyltransferase DPY19 is essential for transmission in *Plasmodium*, but not for asexual blood stage growth so C-mannosylation defects are unlikely to be related to delayed death (25, 26). GPIs function as the membrane anchors for the major surface proteins (GPI-APs) of asexual blood stages as well as being the major class of free glycolipids in the parasite plasma membrane (27, 28). We hypothesise that *P. falciparum* undergoing PRDD have reduced *de novo* synthesis of dolichols and consequently less abundant GPI. Consistent with this hypothesis, we found that apicoplast inhibitors cause surface GPI-APs to lose their membrane association and become untethered in the parasitophorous vacuole. These parasites had defective host cell rupture, consistent with known roles for GPI-anchored proteins in initiating egress. Finally, the cumulative effect of disrupting GPI-APs renders parasites incapable of successful invasion. These data support a crucial role for the apicoplast in generating dolichols needed for synthesis of essential GPIs.

## Results

### Radiolabelling of dolichols and GPI glycolipids in a cell-free system

To quantitate endogenous levels of dolichols as well as downstream products in asexual *P. falciparum* blood stages, we developed a cell-free assay to metabolically label endogenous pools of dolichol phosphate and GPI intermediates. Cell-free extracts (0.5 – 1 x 10□ cell equivalent/mL) were prepared by mechanically lysing trophozoites in a hypotonic lysis buffer, and then labelling with GDP-[^3^H]-mannose (GDP-[^3^H]Man). DPM and GPI glycolipids synthesised from endogenous pools of dolichol-P and GPI precursors were subsequently extracted, recovered by solvent partitioning, and analysed by HPTLC. In the absence of inhibitors, incubation of cell-free extracts with GDP-[^3^H]Man led to the synthesis of a series of glycolipid species that were putatively identified as DPM and GPI intermediates. The latter have previously been shown to contain an acylated PI moiety (three fatty acyl chains total) and either neutral (Man_2-4_-GlcN) or charged (EtN-P-Man_3-4_GlcN) glycan headgroups (29). The identity of these species was confirmed by mild acid treatment (leading to selective loss of labelled DPM), mild base treatment (leading to removal of ester-linked fatty acids and loss of all labelled GPI species) and Jack bean α-mannosidase treatment (leading to selective removal of labelled GPI species containing uncapped neutral mannose backbone, but not ethanolamine-P-capped GPIs or DPM which contains a β-linked mannose) of the labelled extracts (29, 30) (**Fig 2A**).

**Fig 2.**
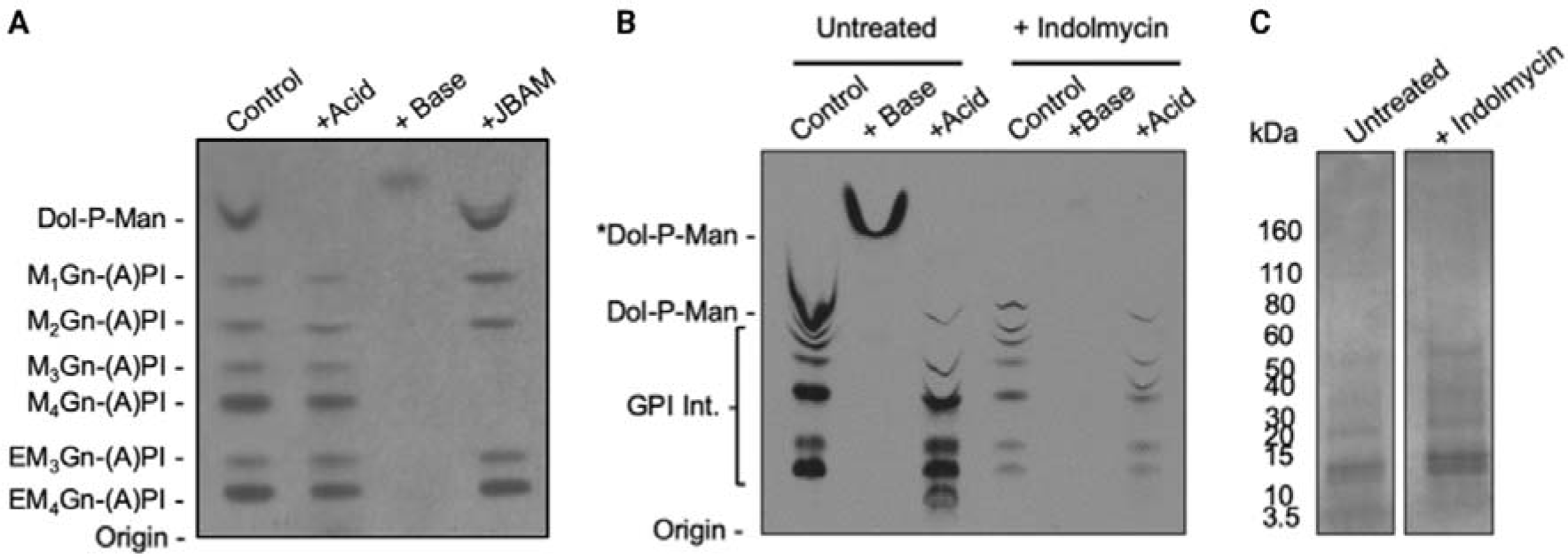
Metabolic labelling of *P. falciparum* glycolipids with GDP-[3H]-mannose. **A)** Fluorograph of HPTLC separating *P. falciparum* ^3^H-Man-labelled GPIs putatively identified according to Gerold et al. 1999 (30). Chemical and enzyme treatments of labelled glycolipids fractions. **B**) Fluorograph of HPTLC separating [^3^H]-Man-labelled *P. falciparum* glycolipids from untreated and indolmycin [50 µM] treated parasites (1x10^9^ cell equivalents/mL) collected 72–78 h post drug administration (equivalent to 28–32 hpi in the second IDC after treatment). The glycolipids were divided into equal fractions and chemically treated as indicated. Dol-P-Man was detected in untreated and was resistant to mild base treatment (+ Base). GPI intermediates were detected in untreated and were resistant to mild acid treatment (+ Acid). In indolmycin treated parasites Dol-P-Man was below the limit of detection and there was less [^3^H]-man incorporation into GPI. **C)** Parasite lysate (1x10^9^ cell equivalents/mL) were analysed by SDS-Page and Coomassie blue as a loading control.

### Dolichol-P-Man pools are depleted during delayed death

We next wanted to determine if isoprenoid depletion during indolmycin-induced delayed death leads to depletion of intracellular pools of dolichol phosphate and *de novo* GPI glycan elongation. Indolmycin is an apicoplast tryptophan tRNA synthetase inhibitor which induces a well characterised delay death phenotype. Cell free extracts of untreated and indolmycin-treated *P. falciparum* trophozoites were metabolically labelled with GDP-[^3^H]Man and extracts analysed by HPTLC. As expected, cell-free extracts of non-drug-treated parasites exhibited high levels of DPM synthesis using endogenous pools of Dol-P, with concomitant labelling of GPI intermediates. Subsequent treatment of the extracts with mild acid or mild base resulted in selective loss of ^3^H-labelled DPM and GPIs, respectively (**Fig 2B**). Note that the HPTLC migration of DPM in both the control and base-treated sample is distorted by the comigration with unlabelled phospholipids and cleaved fatty acids, respectively **(Fig 2B**). In marked contrast, cell-free extracts of indolmycin-treated parasites exhibited greatly reduced incorporation of GDP-[^3^H]Man into DPM and GPI glycolipids. As a control, parasite lysates were analysed by SDS-page and Coomassie blue to confirm equal cell equivalents in these assays (**Fig 2C**). Based on quantitation of ^3^H-labelled GPIs and Dol-P-Man in the HPTLC, delayed death parasites had a 30-fold and 3-fold reduction in labelling for Dol-P-Man and GPIs, respectively (**S1 Fig**) (31). Collectively, these results suggest that indolmycin treatment leads to delayed death which is associated with a dramatic decrease in intracellular pools of Dol-P and DPM, with a concomitant decrease in the rate of GPI biosynthesis

### GPI-anchored proteins MSP1 and MSP2 mislocalise when treated with delayed death drugs

GPI-APs contain an N-terminal signal sequence, which is required for protein transit into the ER lumen, and a short C-terminal hydrophobic sequence, which initially tethers the protein to the inner leaflet of the ER. The C-terminus is rapidly cleaved by the multi-protein GPI-transamidase complex at the ω-site allowing the formation of an amide bond between the newly-created C-terminus of the target protein and the ethanolamine phosphate of the pre-synthesised GPI (32). Much of the GPI-anchored proteome of the IDC is expressed in the later trophozoite and schizont stages and is localised to the parasite surface where GPI-anchored proteins (GPI-APs) constitute >60% of the total surface protein coat (33). When the GPI biosynthetic pathway in other eukaryotes is impaired either via mutations in critical biosynthetic enzymes or chemical inhibition, normal localisation of GPI-anchored proteins is disrupted (34). In the absence of GPI precursors, GPI-APs can accumulate within the ER, leading to induction of an ER stress response as well as their retrotranslocation across the ER membrane via ER-associated protein degradation (35, 36). Alternatively, non-anchored proteins may enter the secretory pathway and be exported into the extracellular environment (37). To determine the fate of GPI-APs following the loss of apicoplast function, PRDD was induced with clindamycin (an inhibitor of the apicoplast 50S ribosomal subunit), and simultaneously rescued with the prenylation precursor GGOH. Parasites were then harvested at the second cycle schizont stage and trafficking of the two most abundant merozoite surface proteins, MSP1 and MSP2 (comprising >50% of the merozoite surface coat) monitored by immunofluorescence microscopy (**Fig 3A**) (33). MSP1 and MSP2 were both trafficked to the plasma membrane of newly formed merozoites towards the end of schizogony (**Fig 3B**), and localised adjacent to the non-GPI microneme membrane protein, apical membrane antigen 1 (AMA1) (**Fig 3C**). Importantly, both proteins exhibited distinct distribution from Exported Protein 2 (EXP2) which forms nutrient/protein transport pores in the PVM (**Fig 3D**). In marked contrast, the localisation of both GPI-APs was perturbed in PRDD schizonts, a phenotype that was also observed when PRDD was induced with indolmycin (**S2 Fig**). Expression of both proteins was reduced, and residual fluorescence was associated with puncta that overlapped with EXP2, indicating a PVM lumen/membrane localisation. This was not associated with a global defect in secretory protein transport, as the localisation of AMA1 (and EXP2) was unaffected (**Fig 3C,D**). Unlike typical delayed death where IMC integrity is lost following failure to prenylate Rab11a, IMC protein glideosome associated protein 45 (GAP45) showed a typical IMC localisation after clindamycin treatment (**Fig 3E**) (14).

**Fig 3.**
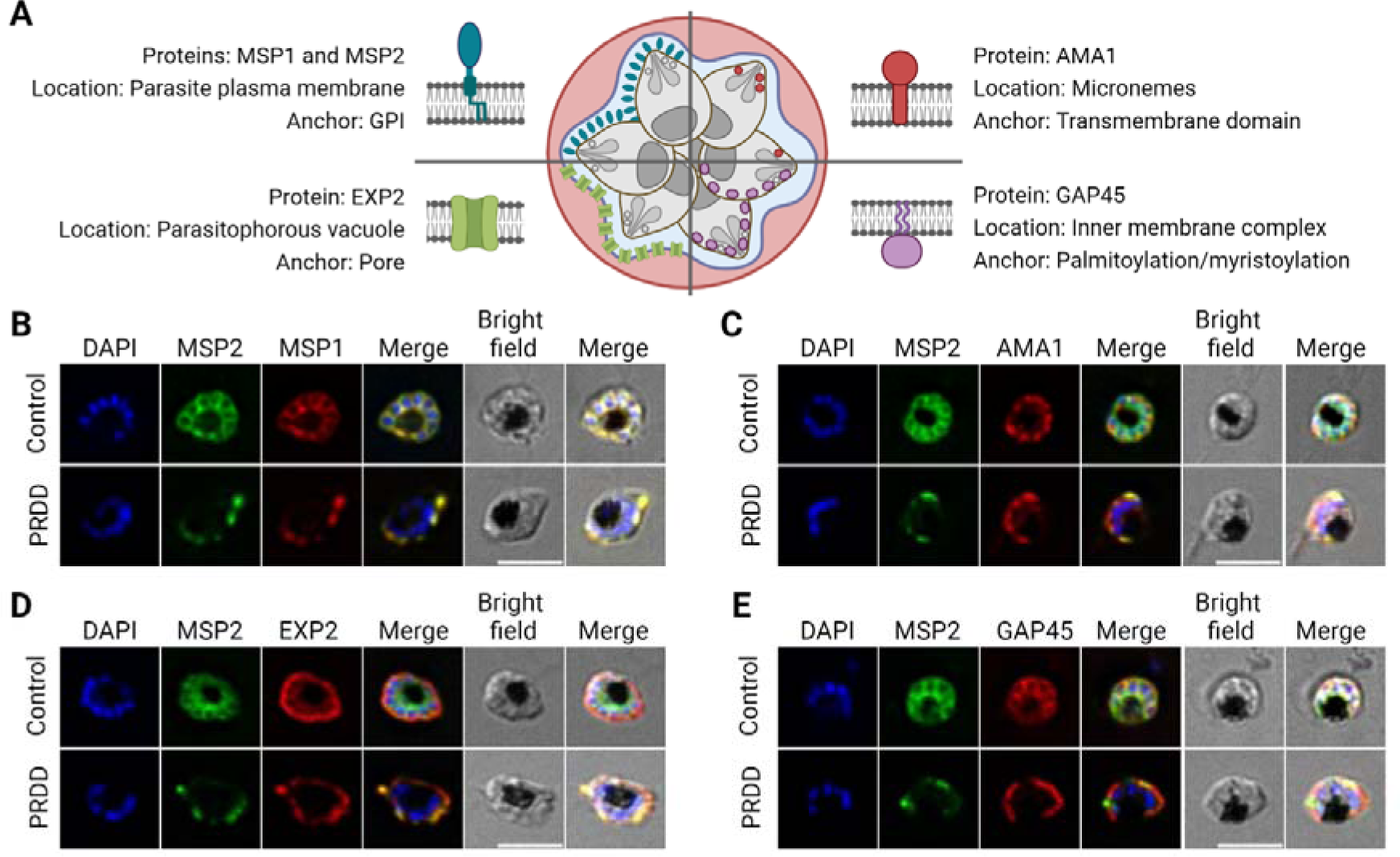
Surface GPI-anchored proteins lose their membrane association in second cycle partially rescued delayed death (PRDD) *P. falciparum*. **A)** The localisation of surface GPI-anchored proteins was examined along with proteins associated with membranes via dolichol independent mechanisms. The GPI-anchored protein merozoite surface protein 2 (MSP2) (green) was co-localised (red) with **B)** GPI-anchored MSP1 **C)** transmembrane protein apical membrane antigen (AMA1), **D)** membrane spanning pore exported protein 2 (EXP2) and **E)** glideosome associated protein 45 (GAP45) – membrane associated by attachment of palmityl and myristoyl groups. Apicoplast inhibition with translation inhibitors results in surface GPI-anchored proteins lose their membrane association. Nuclei stained with 4,6-diamidino-2-phenylindole (DAPI; blue). Scale bar = 5 µm.

### Localisation of surface GPI-APs is not affected by fosmidomycin treatment

Direct inhibition of IPP biosynthesis with the DOXP reducto-isomerase inhibitor fosmidomycin results in cell death within the same cycle during which it is administered, killing parasites by the trophozoite stage, rather than delayed death (8). This is likely mediated by the loss of function of prenylated Rab-mediated haemoglobin trafficking to the digestive vacuole, which is essential for trophozoite development (18). Given that protein prenylation is adversely affected by fosmidomycin treatment, we tested whether GPI-AP localisation is also sensitive to fosmidomycin. Consistent with previous reports, GGOH supplementation allowed fosmidomycin treated parasites to mature to schizont stages (**Supp Fig 3A**). These parasites showed normal IMC formation and MSP2 localisation, indicative of maintained prenylation and GPI biosynthesis during this first cycle and contrasting with the phenotype seen in the second cycle of delayed death. Continued GPI biosynthesis and protein anchoring following fosmidomycin treatment suggests there is an initial, residual pool of dolichols that supports GPI biosynthesis, consistent with previous reports that low levels of dolichols are sufficient to maintain glycosylation (23). As dolichols are not consumed through their participation in GPI biosynthesis, they appear to be more robust in the short term to fluctuations in IPP (**S3 Fig**). In contrast, prenyl groups are covalently linked to the C-terminus of proteins with prenylation motifs and thereby consumed, so are dependent on the *de novo* synthesis of IPP for continuing prenylation.

### PRDD leads to surface GPI-anchored proteins losing their membrane attachment

The mislocalisation of MSP1 and MSP2 in PRDD parasites shown by IFA (**Fig 3B**) is consistent with a reduction in available mature GPIs leading to MSP1 and MSP2 losing their membrane tethers. To biochemically test this, we performed a cell fractionation detergent solubility assay (**Fig 4A**). Soluble proteins in the red blood cell (RBC) and parasitophorous vacuole (PV) were collected using saponin, a detergent that at low concentrations selectively lyses the RBC and PV membranes while leaving the parasite itself intact. The parasite material was subsequently extracted with Triton X-114 (TX-114), and phase partitioned to separate GPI-AP that retained (detergent phase) or lacked (aqueous phase) a GPI anchor.

**Fig 4.**
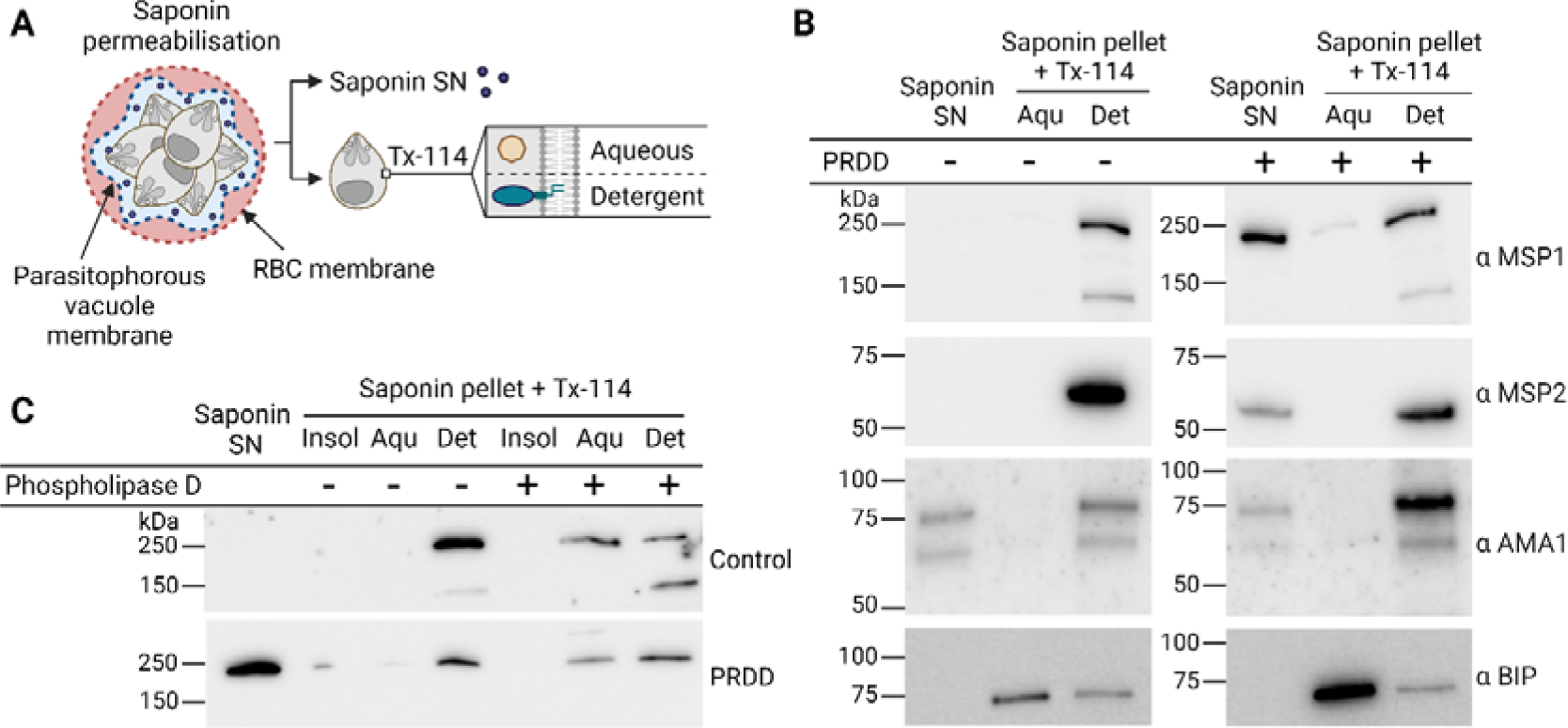
Detergent solubility assays further confirm the PV localisation of GPI-anchored proteins following drug treatment. **A)** Following permeabilisation by saponin and centrifugation, the resulting supernatant (SN) is collected to assay for proteins soluble in the red blood cell (RBC) or parasitophorous vacuole (PV). The parasite pellet is then treated with Triton X-114 (TX-114) and phase separation is used to partition proteins into an aqueous and detergent phase to enrich for soluble and membrane associating proteins respectively. **B)** Representative western blots showing that GPI-anchored proteins MSP1 and MSP2 partition in the detergent phase, with AMA1 and BIP as the as the membrane bound and soluble control respectively. After apicoplast inhibition MSP1 and MSP2 are found in the saponin supernatant, consistent with a portion of these proteins having lost their GPI and releasing into the PV. **C)** Treatment of parasite extracts with a GPI-specific phospholipase D removes the membrane anchor of MSP1, causing it to become enriched in the aqueous phase.

Both MSP1 and MSP2 were found in the soluble, saponin fraction of PRDD parasites but not the no-drug control, consistent with the IFA data showing that after apicoplast inhibition, a portion of these proteins lose their membrane attachment and become soluble within the PV (**Fig 4B**). While MSP2 has a predicted molecular weight of ∼28 kDa, we consistently detected migration with an apparent molecular weight higher than this size with multiple independent monoclonal primary antibodies, as previously reported for independent monoclonal and polyclonal reagents (**S4 Fig**) (38, 39). AMA1, which is membrane-associated through its transmembrane domain rather than a GPI-anchor, is unaffected by apicoplast inhibition, consistent with the changes in MSP1 and MSP2 being due to GPI-anchor loss. AMA1 is detected in both full length and processed forms (83 and 66 kDa), a sign of schizont maturation (**Fig 4B**).

A pool of MSP1 and MSP2 is observed in the detergent fraction of PRDD parasites, indicating some residual membrane association. This may represent 1) normal GPI-AP at the surface, 2) proteins with a native hydrophobic C-terminal sequence being retained in the ER without processing by the transamidase or 3) proteins without an anchor themselves, but strong association to a membrane bound protein, as seen with ER chaperone BIP. To probe this further, saponin-lysed parasite extracts were incubated with the GPI-specific phospholipase D to cleave the GPI anchor and liberate their associated proteins into the aqueous phase (**Fig 4C**). The sensitivity of residual MSP1 to the GPI-specific phospholipase during PRDD suggests these parasites retain a minor capacity to synthesise or recycle GPI-anchors.

### Depletion of dolichols by apicoplast inhibition culminates in a second cycle lethal egress defect

PRDD parasites form schizonts which have abnormal morphology and are unable to establish a third intraerythrocytic development cycle (14) (**Fig 5A**). To investigate whether PRDD parasites can egress, we used a transgenic *P. falciparum* 3D7 line expressing a NanoLuc luciferase (Nluc) which is trafficked into the host RBC cytoplasm (40). When the RBC is ruptured during normal egress, the Nluc enzyme is released into the supernatant and Nluc activity can be assayed following the addition of the NanoGlo substrate (**Fig 5B**). PRDD and control schizonts were purified and incubated with either compound 1 (C1, at 4 µM), a 3’5’-cyclic monophosphate (cGMP)-dependent protein kinase inhibitor that reversibly blocks egress or DMSO vehicle. PRDD parasites showed comparable Nluc activity to their C1 controls (22.62% vs 22.28% at 4 hours and 42.90% vs 37.21% at 8 hours), indicating that these parasites have a severe egress block. The continued egress inhibition over the 8-hour incubation indicates that this phenotype is not merely due to drug induced cell cycle delays, but a lasting biological impairment (**Fig 5C**).

**Fig 5.**
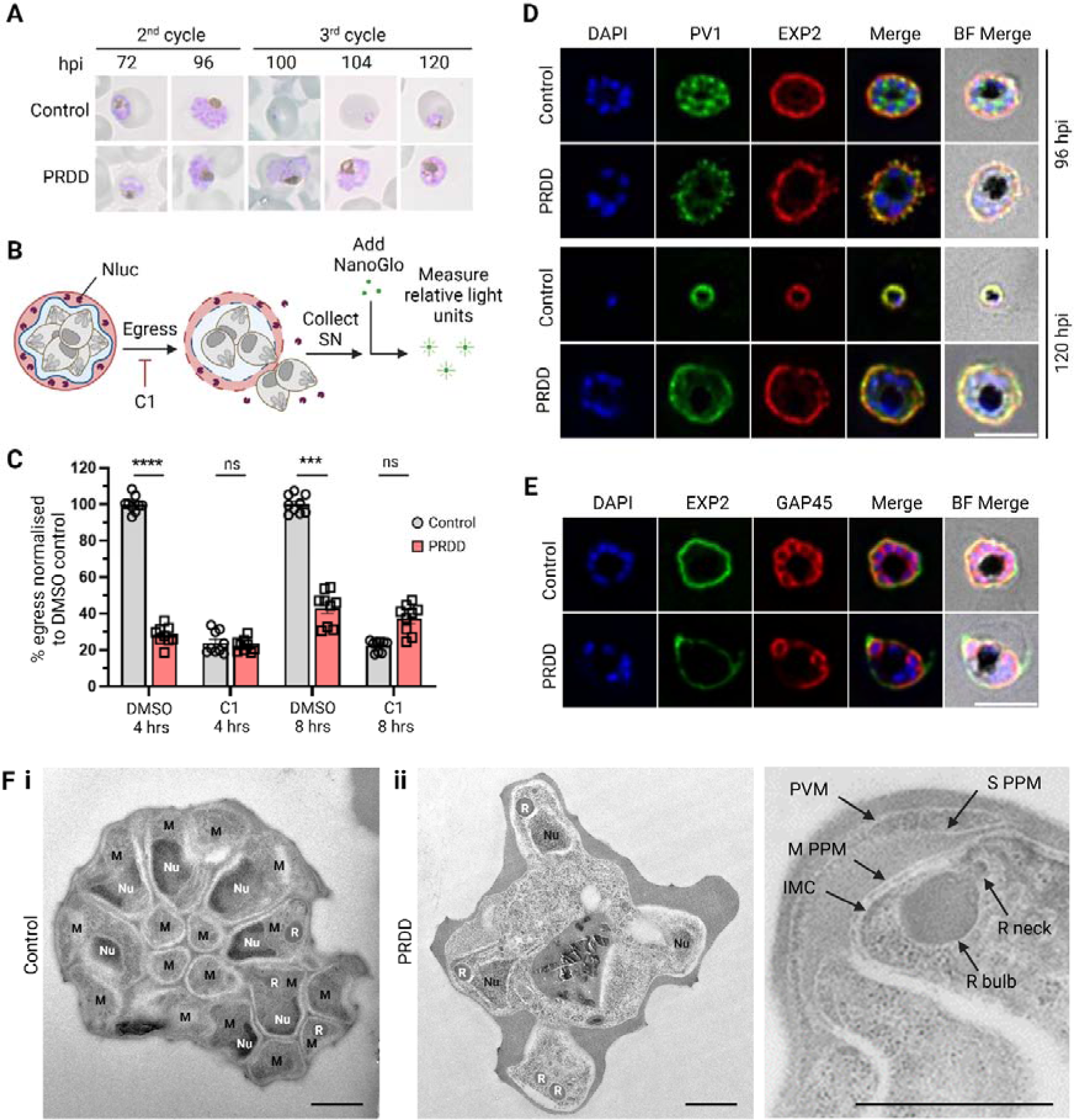
Second cycle PRDD schizonts cannot efficiently segment or egress from their host RBC. **A)** Giemsa blood smears show that PRDD stall at the schizont stage and are unable to establish a third cycle of infection. **B)** To test parasites egress, an assay was performed with parasites transfected with an exported Nanoluciferase (Nluc). Upon RBC lysis, Nluc is released into the supernatant, which is collected. The NanoGlo substrate is added, with the relative light released used as a proxy for egress. Parasite egress can be reversibly blocked with the inhibitor compound 1 (C1). **C)** Purified schizonts were plated out in the presence of either DMSO vehicle or C1 at 4 µM and the supernatant collected after 4 or 8 hours. PRDD parasites showed egress comparable to incubations with C1. Each dot represents a technical replicate normalised to the DMSO 100% control, with 3 technical replicates in each of the 3 biological experiments. Error bars = SD. Unpaired t-test, p value < 0.05. **D)** IFAs showing persistence of the parasitophorous vacuole membrane (PVM) structure as seen by maintained EXP2 labelling. The egress defect of PRDD schizonts involves an inability to completely rupture their PVM long after typical egress occurs. BF, bright field **E)** These schizonts are also unable to segment appropriately, with new inner membrane complex (IMC) forming around only some nuclei as seen by labelling of the IMC protein GAP45. Nuclei stained with DAPI (blue). Images represent single Z stacks. Scale bar = 5 µm. **F)** Electron microscopy shows that while some nuclei (Nu) are not successfully segregated and enveloped by a new parasite plasma membrane (PPM), apparently normal rhoptry formation still occurs in PRDD parasites. Hpi, hours post invasion; M, merozoite; R, rhoptry; S PPM, schizont parasite plasma membrane; M PPM, merozoite parasite plasma membrane. Scale bar = 1 µm

We next aimed to discern the stage of egress arrest observed in the PRDD parasites. For successful egress, parasites initiate the poration and then rupture of first the PV membrane and then the erythrocyte membrane. The only known GPI-anchored protein implicated in egress of the asexual stages is MSP1. Following activation by the protease SUB1, MSP1 binds the host cell cytoskeleton protein spectrin (41). This interaction inflicts significant mechanical stress which is necessary for efficient RBC rupture. Using a transgenic parasite line expressing 3xHA-tagged parasitophorous vacuolar protein 1 (PV1-HA) we performed IFAs on parasites 96- and 120-hours post invasion. PRDD schizonts appeared to partially perforate the PVM, as shown by the fragmented and discontinuous EXP2 labelling and the leakage of PV1-HA and EXP2 into the erythrocyte cytosol (**Fig 5D**). The nuclei remain restricted within the PVM boundaries, suggesting that the PVM perforates but does not fully rupture. Mechanistically this egress defect is not explained by our current understanding of MSP1 loss-of-function phenotype, as the MSP1-spectrin interaction is only possible following PVM rupture. These data therefore support disruption of a GPI-anchored protein other than MSP1, although no known GPI-APs have yet been implicated in PVM rupture.

### Partially rescued delayed death schizonts do not segment properly

During segmentation, the parasite plasma membrane (PPM) invaginates around each nuclei to form discrete merozoites within the PV. A reduction in inter-merozoite PV1-HA in PRDD parasites (**Fig 5D**) suggests membrane partitioning of individual merozoites is compromised. IFAs probing an IMC marker (GAP45) showed that PRDD parasites retain their original IMC but were unable to efficiently form new IMCs around all daughter merozoites (**Fig 5E**). To further elucidate this segmentation defect, we performed electron microscopy (**Fig 5F**). Compared to control samples, the PRDD parasites produced notably fewer merozoites with some daughter nuclei lacking encapsulation by new membrane architecture such as an IMC and PPM, although some parasites contained normally segmented merozoites (**S5 Fig**). The *Plasmodium* GPI-anchored protein rhoptry-associated membrane antigen (RAMA) is essential for rhoptry neck formation, however rhoptries with distinct bulb and necks were detected in PRDD merozoites.

### Merozoites from PRDD are unable to invade new RBCs

While PRDD *P. falciparum* has an egress defect during *in vitro* culturing conditions such that we see no reinvasion after the second cycle, it is possible that the mechanical stress experienced in the bloodstream of an *in vivo* infection may be sufficient to rupture the host cell and allow daughter merozoites to escape. Merozoite surface GPI-APs are implicated in facilitating weak binding with ligands at the RBC surface necessary for efficient invasion.

To determine whether merozoites derived from PRDD parasites are capable of invasion, a flow cytometry-based invasion assay was performed using merozoites mechanically released from schizonts (**Fig 6A**). Late stage second cycle schizonts were magnet-enriched and further matured at least 4 hours in the presence of egress inhibitor trans-Epoxysuccinyl- L-leucylamido (4-guanidino) butane (E64). Daughter merozoites were then mechanically purified using a 1.2 µm syringe filter. Merozoite viability was confirmed using live cell microscopy where parasites were co-stained with the DNA marker Hoechst 33342 and MitoTracker™, a membrane potential dye which only stains live cells (**Fig 6B**). Merozoites were then introduced to uninfected RBCs with GGOH and allowed to reinvade in the presence or absence of invasion inhibitor heparin. After 20 h, parasitaemia was determined by flow cytometry (**Fig 6C**).

**Fig 6.**
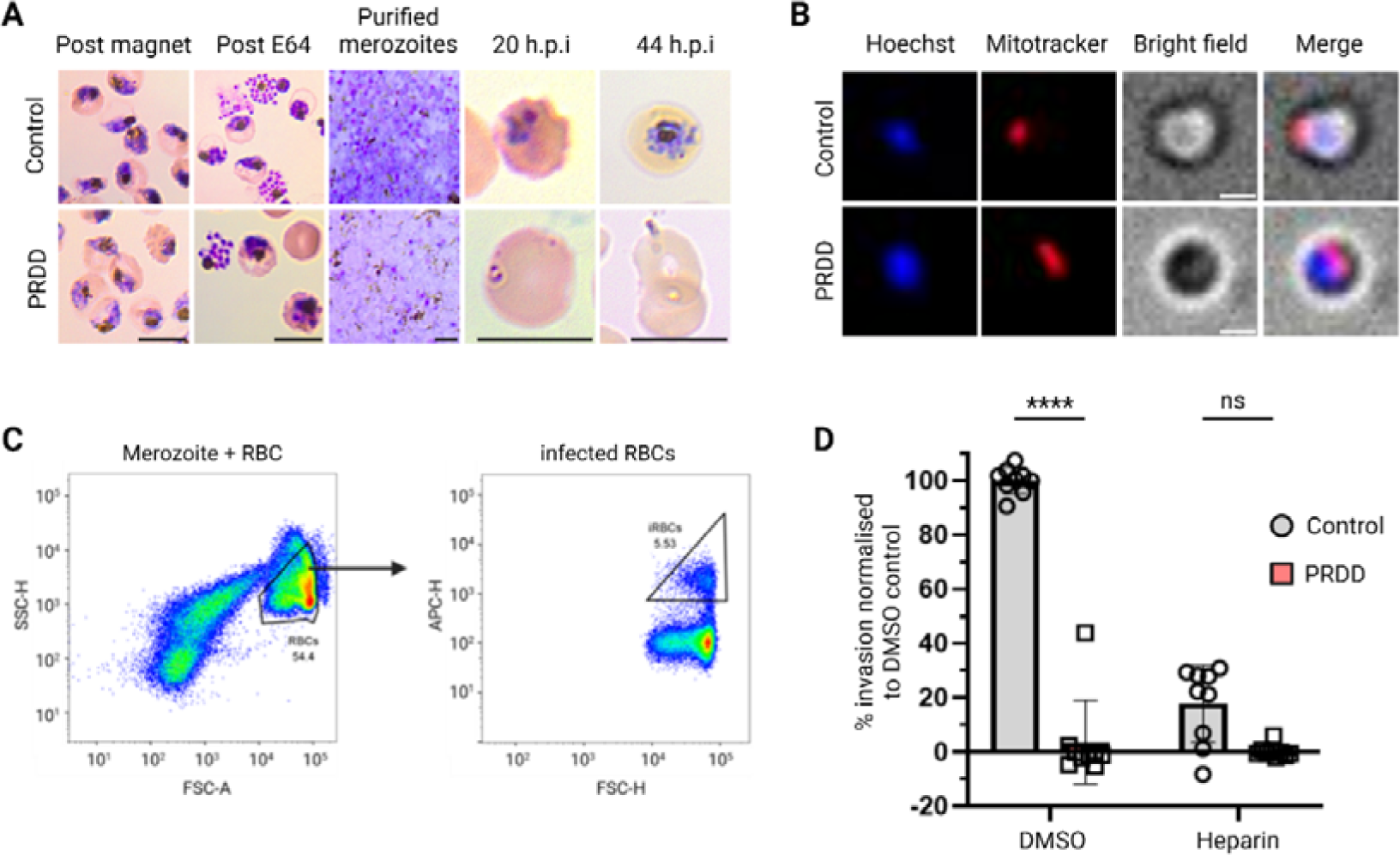
Merozoites derived from PRDD second cycle schizonts are incapable of invasion. **A)** Giemsa smears show parasite development. Magnet enriched schizonts matured in the presence of E64 before their merozoites were mechanically extracted from the host RBC. Merozoites were incubated with fresh RBCs and parasite growth was monitored over 20 and 44 hours. Scale bar = 7 µm. **B)** Live cell imaging of purified merozoites with MitoTracker™ staining shows parasites are viable following their purification. To determine whether merozoite have invaded, flow cytometry was performed 20 hours after parasites were added to fresh RBCs. Scale bar = 1 µm. **C)** Gate setting for flow cytometry analysis performed using FLowJo software. RBCs were selected using SSC-H and FSC-A gating to exclude uninvaded parasites. **D)** PRDD parasites have negligible invasion, comparable to that of parasites treated with the invasion inhibitor heparin. Each dot represents a technical replicate normalised to the DMSO 100% control, with 3 technical replicates in each of the 3 biological experiments. Error bars = SD. Unpaired t-test, p value < 0.05.

Despite loading equal numbers of schizonts, more merozoites were recovered from the filtration of the control parasites compared to PRDD, as expected given that segmentation was impaired following apicoplast inhibition. Even accounting for this merozoite loading factor, parasites with reduced surface GPI-APs were incapable of invasion (**Fig 6D**), with Giemsa smears suggesting that PRDD derived merozoites do attach to the surface of RBCs but fail to penetrate the host cell (**Fig 6A**). Together these data suggests that the collective loss of surface GPI-APs results in the complete ablation of invasion.

## Materials and methods

### P. *falciparum* cell culture

Asexual blood stage *P. falciparum* was cultured as previously described in O positive human red blood cells (RBCs) (Australian Red Cross Lifeblood) in RPMI 1640 with HEPES (Formedium) supplemented with 0.25% (w/v) Albumax II (Gibco), 0.5 mM hypoxanthine and 42 μM gentamicin (Santa Cruz), referred to as media. Cultures were maintained at 37 °C in an atmosphere of 1% O□, 5% CO□, 94% N□.The transgenic Nluc parasite line (PEXEL-Nluc), produced by the transfection of 3D7 *P. falciparum* with the Hyp1-Nluc plasmid, was a kind gift from Paul Gilson (Burnet Institute, Melbourne, Australia) (40). Transgenic parasites were maintained by culturing with 2.5 nM WR99210 (Jacobus, USA). The transgenic parasite line PV1-HA was also maintained by culturing with 2.5 nM WR99210 (Jacobus, USA) (42).

### *P. falciparum* synchronisation

Synchronised *P. falciparum* cultures was achieved by treatment with 5% (w/v) D-sorbitol at the ring stage for at least two consecutive cycles prior to experimentation. When tight synchronisation was desired, schizont stage parasites were enriched via 65% (v/v) Percoll cushion, incubated for 4 hours with fresh RBCs, and then treated with 5% (w/v) D-sorbitol.

### 3H-Mannose labelling of dolichol phosphate mannose and GPIs in a cell-free assay

Samples were prepared for cell-free assay (methods developed from (29, 30, 43)) using synchronised 0–4 hours post invasion (hpi) ring-stage parasites treated with 50 μM indolmycin or DMSO vehicle control for 30 h. Media was replaced 30 h post drug administration. Trophozoites (5–10% parasitaemia) were collected at approximately 72–78 h post drug administration (equivalent to 28–34 hpi in the second IDC) and isolated by 0.05% (w/v) saponin treatment, pelleted by centrifugation and resuspended in ice-cold PBS.

Trophozoites were divided into equal aliquots with 5 × 10^7^–1 × 10^8^ cells per assay. All steps were carried out on ice or at 4 °C. The parasites were pelleted by centrifugation at 25,000 x g and resuspended in 5 × the pellet volume with hypotonic lysis buffer (1 mM HEPES-NaOH pH 7.4, 2 mM EGTA, 2 mM DTT with protease inhibitor cocktail; cOmplete™, EDTA-free, Roche). The parasite suspension was incubated for 10 min on ice and then lysed by approximately 30 passages through a 1mL syringe fitted with a 27G needle.

Equal volumes of 2 × assay buffer (200 mM HEPES-NaOH pH 7.4, 100 mM KCl, 10 mM MgCl_2_, 10 mM MnCl_2_, 4 mM EGTA, 4 mM DTT with protease inhibitor cocktail) were added to each sample and the lysed cells were pelleted by centrifugation (15,000 x *g*, 1 min). The supernatants were aspirated, the pellets washed once in 60 μL with 1 × assay buffer, then resuspended in 100 μL 1 × assay buffer (containing GDP-[^3^H]-mannose (GDP-[^3^H]Man, 2E+05 cpm).

The reactions were incubated for 1 h at 37 °C and terminated by addition of chloroform: methanol (1:2, v/v) giving a final monophasic ratio of chloroform: methanol: aqueous (1:2:0.8, v/v). Glycolipids were then extracted, with intermittent bath sonication, for 18h at 4 °C and the supernatants (lipid extracts) were transferred to fresh microfuge tubes and dried under N_2_ gas prior to 1-butanol partitioning.

### 1-Butanol aqueous partitioning

The glycolipids extracted from control and drug-treated parasites were recovered by partitioning in a biphasic system of 1-butanol (200 μL) and water (150 μL). After centrifugation, the upper butanol phase was removed, and the lower aqueous phase washed once with water-saturated 1-butanol (200 μL). The combined butanol phases were then back-extracted with water (150 μL) and dried in a centrifuge vacuum concentrator. Glycolipids recovered in the 1-butanol phase were resuspended in 40% (v/v) 1-propanol and aliquots taken for scintillation counting prior to chemical and enzyme treatments or high-performance thin layer chromatography (HPTLC) analysis.

### HPTLC analysis of glycolipids

Samples were resuspended in 6 μL 40% (v/v) 1-propanol with vortexing and bath sonication were applied to Silica Gel 60 aluminium-backed HPTLC plates (MERCK) (30, 44). HPTLC plates were developed for 10 cm in chloroform, methanol, 13M NH_4_OH, 1M ammonium acetate, water (180:140:9:9:23, v/v). The radioactive lanes were scanned using a Berthold LB2821 automatic TLC linear analyser prior to fluorography which was performed by spraying with En^3^Hance™ (DuPont) and exposing the HPTLC to BioMax® Maximum Resolution (MR) Autoradiography Film (Kodak) at -80 °C.

### Chemical and enzyme treatments

Mild acid hydrolysis of the *P. falciparum* glycolipids was performed in 0.5 M HCl in 50% (v/v) 1-propanol (50 µL) for 15 min at 55 °C. Mild base hydrolysis was performed in 0.1 M methanolic-NaOH (50 µL) for 2 hr at 37 °C and samples neutralised with 1 M acetic acid. Digestion with Jack bean α-mannosidase (50 units/mL) was performed in 0.1 M sodium acetate, pH 5.0 containing 0.2% taurodeoxycholic acid (25 µL) for 18hr at 37 °C. In each case, the lipidic products were recovered by 1-butanol: water biphasic partitioning as described above. The dried 1-butanol phases were resuspended in 6 μL 40% 1-propanol for analysis by HPTLC and fluorography (see HPTLC analysis of glycolipids).

### Induction of partial rescue delayed death (PRDD)

Synchronised early ring stage *P. falciparum* cultures (4-8 hpi) were treated with apicoplast inhibitors (either 5 µM clindamycin (Sigma-Aldrich, PHR 1159) for > 50 times the delayed death IC₅₀, or 50 µM indolmycin (BioAustralis Fine Chemicals) for ∼ 50 times the delayed death IC₅₀) and supplemented with 5 µM GGOH (Sigma Aldrich, G3278) to induce delayed death while rescuing the prenylation defect. Control cultures were supplemented with 5 µM GGOH. Spent medium was replaced and re-supplemented with 5 µM GGOH in the second intraerythrocytic development cycle.

### Indirect Immunofluorescence assays

Glass coverslips were pre-treated with 0.1 mg/mL pHAE (erythroagglutinating phytohemagglutinin) (Sigma Aldrich) and infected RBCs at 5% haematocrit in phosphate buffered saline (PBS) were coated as a monolayer. Cells were then fixed with 2% (v/v) paraformaldehyde/0.008% (v/v) glutaraldehyde for 20 minutes, permeabilised in 0.1% (v/v) Triton X-100 (TX-100) for 10 minutes then blocked with 3% (w/v) bovine serum albumin (BSA) for 1 hour. Primary antibodies: 0.04 µg/mL mouse anti-MSP2 (clone 9D11) (39), 1:900 rabbit anti-MSP1 (R644), 1:1000 rabbit anti-GAP45 (45), 1:500 rabbit anti-AMA1 (46), 1:1000 rabbit anti-EXP2 (47), 1:300 rat anti-HA (Sigma Aldrich, clone 3F10) in 3% (w/v) BSA were incubated for 2 hours. Secondary antibodies: 1:300 goat anti-mouse Alexa Fluor® 488 (Abcam, ab150117), 1:300 goat anti-rabbit Alexa Fluor® 594 (Invitrogen, A11012) and 1:300 donkey anti-rat Alexa Fluor® 488 (Abcam, ab150156) in 3% (w/v) BSA were added for 1 hour before addition of 300 nM DAPI (Sigma Aldrich) and ProLong® Gold Antifade (Cell Signaling Technology). Images were captured and deconvolved using the DeltaVision Elite Widefield (GE Healthcare) platform. Brightness and contrast adjustments and image merging was performed using ImageJ (version 1.53) (48).

### TX-114 phase separation

Synchronised *P. falciparum* cultures were treated with vehicle or had PRDD induced at the ring stage. Approximately 90 hours after treatment schizont infected red blood cells were isolated using a 65% Percoll cushion and washed twice in PBS. To permeabilise the RBC and PVM, infected red blood cells (iRBCs) were then lysed in 0.03% w/v saponin for 10 minutes on ice. Following centrifugation, the resulting saponin supernatant containing RBC and PV soluble proteins was retained and the parasite pellet was washed three times in PBS before solubilisation in 5% (v/v) Triton X-114 (TX-114) (Sigma-Aldrich) (0 °C, 30 min) that had been precondensed as previously described (49). Samples were centrifuged at 4 °C and the cleared supernatant was laid over a 6% (v/v) sucrose cushion and incubated for 5 minutes at 30 °C to achieve the TX-114 cloud point and phase-partitioning. The aqueous and detergent fractions were separated by centrifugation, followed by a second round of TX-114 treatment and phase-partitioning of the aqueous fraction. Proteins were precipitated from the TX-114 aqueous and detergent phases by adding trichloroacetic acid (TCA) to a final concentration of 13% (v/v), and the protein pellets washed twice in ice cold acetone.

### PI-PLD digest

The second cycle PRDD and control schizonts were harvested via Percoll and lysed with saponin as described above. Parasites were resuspended in 0.1% (v/v) TX-100 (Sigma) in PBS, 2.5 mM CaCl□, protease inhibitor cocktail and solubilised on ice for 30 minutes. The samples were then treated with human serum containing phospholipase D (Australian Red Cross Lifeblood) or 0.25% (w/v) Albumax II (Gibco) and incubated for 3 hours at 37 °C. The samples were then chilled on ice and diluted with 5% (v/v) TX-114 to achieve a final concentration of 4% (v/v) TX-114 and 0.02% (v/v) TX-100. Samples were centrifuged at 4 °C and the cleared supernatant phase-partitioned over a 6% (v/v) sucrose cushion as described above.

### Western blotting

Lysates were resuspended in sample buffer (Laemmli sample buffer with β-mercaptoethanol, BioRad) and boiled at 90 °C for 10 minutes. Proteins were separated by size using SDS PAGE 4-15% gels (BioRad) at 160 V, 50 minutes and then transferred to a nitrocellulose membrane using an iBlot2 (Thermo Fisher). Membranes were blocked in 5% (w/v) skim milk in PBS for 1 hour at room temperature on an orbital shaker before incubating with primary antibodies rabbit anti-MSP1 (R644) 1:1,000, mouse anti-MSP2 (9D11, 11E1, 2F2) 1 µg/mL (39), rabbit anti-AMA1 1:1,000 and mouse anti-BIP 1:1,000 in 3.5% (w/v) skim milk overnight at 4 °C. Membranes were washed in 0.05% (v/v) TWEEN20 in PBS (PBST) before incubating with the secondary antibodies goat anti-mouse HRP, 1:10000 (Promega: W4021) and in 3.5% (w/v) skim milk for 1 hour at room temperature. Blots were again washed in PBST with a final rinse in PBS. Membranes were developed (Clarity™ peroxide reagent and luminol/enhancer reagent, BioRad 1705061) as per manufacturer’s instructions and imaged using a BioImager (BioRad).

### Egress assay

Nluc parasites were tightly synchronised as described above and PRDD was induced. Approximately 94 hours later, second cycle schizonts were purified using a 65% (v/v) Percoll cushion and adjusted to ∼2% parasitemia and 1% haematocrit in a 96 well plate. Compound 1 (C1, 4 µM) or DMSO was administered in triplicate. Plates were incubated at 37 °C for either 4 or 8 hours, following which the media supernatant containing released NanoLuciferase was collected.

### Measuring NanoLuciferase activity

NanoLuciferase activity was measured as previously described (50). Briefly, 5 µL of the schizont culture was aliquoted into white 96 well luminometer plates to which 45 µL of 1x NanoGlo Lysis buffer with 1:1000 NanoGlo substrate (Promega, USA) was added. Initial parasitemia was determined by measuring the total Nanoluciferase release from RBCs, allowing normalisation between samples. Relative light units (RLU) were measured using the FLUOstar Omega.

To measure parasite egress, the media supernatant was removed at either 4 or 8 hours and 5 µL of supernatant was added to a white 96 well luminometer plate. 45 µL of 1x NanoGlo Lysis buffer with 1:1000 NanoGlo substrate (Promega, USA) was added. RLUs were measured using FLUOstar Omega and Nluc activity was normalised to the initial parasitemia and relative to the no drug control (100% Nluc activity) for the 4- or 8-hour time point.

### Merozoite purification

Tightly synchronised ring stage parasites had PRDD induced (final 4% parasitaemia, 120 mL culture, 2% haematocrit) then schizonts were magnet purified (as described above) between 92-96 hours post drug treatment (51). The enriched schizonts were returned to culture and incubated for 4 hours with an egress inhibitor, E64 (10 μM) (E3132, Sigma-Aldrich), to ensure merozoites have fully developed. After E64 treatment, schizonts were pelleted by centrifugation (1900 x *g*, 7 minutes) and the supernatant was discarded. The schizont pellet was resuspended in 1 mL of media and then immediately filtered through a 1.2 µm, 32 mm Acrodisc® Syringe Filter (Pall Life Sciences) to mechanically rupture the iRBCs and release the daughter merozoites.

### Live cell microscopy of merozoites

Second cycle schizonts were purified and matured in the presence of E64 as described above. Cultures were centrifuged to remove the E64 and schizonts were resuspended in 1 mL of pre-warmed media with MitoTracker™ Red CMXRos (10 nM) (M7512, ThermoFisher). These cells were incubated for 30 minutes at 37 °C, then centrifuged and the supernatant was removed. Schizont pellet was resuspended in 1 mL of media and stained with Hoechst 33342 (1 µg/mL) and then immediately mechanically egressed using a 1.2 µm, 32 mm Acrodisc® Syringe Filter (Pall Life Sciences). The merozoite pellet was washed in media before being resuspended in 2 µL of media, then transferred onto a coverslip and mounted on a glass slide. Samples were imaged with a DeltaVision Elite restorative widefield deconvolution imaging system (GE Healthcare). Deconvolved images were processed using the FIJI ImageJ software (48).

### Electron microscopy

3D7 parasites had PRDD induced, and were magnet purified at 94-98 hpi. iRBCs were fixed in 1% (v/v) glutaraldehyde in RPMI-HEPES, infiltrated with LRWhite resin (ProSciTech) and polymerised with LRWhite chemical accelerant. Ultrathin 100 nm sections of the resin stub were cut using a Leica UC7 ultramicrotome, stained with Aqueous uranyl acetate and Reynolds lead citrate before observation on a Tecnai F20 Transmission electron microscope. Sections were imaged at 120 kV.

### Flow cytometry-based invasion assay

Filter extracted merozoites were added to uninfected RBCs at 1% haematocrit in media supplemented with 5 µM GGOH (Sigma Aldrich, G3278) in the presence or absence of the invasion inhibitor heparin (100 mg/mL) (H3393, Sigma-Aldrich). For relative quantification of merozoites loading, filtered merozoites were added to wells lacking RBCs. Plates were incubated on a shaking incubator (300 rpm, 30 minutes) to allow for invasion events before returning to static culture.

Flow cytometry was performed 20-24 hours later to measure successful invasion of the purified merozoites. iRBCs were washed twice with media to remove extracellular merozoites before being aliquoted in triplicate into a 96-well U-bottomed plate and stained with the nucleic acid dye SYTO-61 (2 μM) (Thermo Fisher) for 15 minutes. PBS was added to each sample to cease the reaction and allowed to incubate for 30 minutes before flow cytometric analysis. Flow cytometry was performed using a FACSCanto II (BD Biosciences). To detect parasites (SYTO-61), samples were excited with the allophycocyanin (APC) (660/20 nm) filter set. Data were analysed using FlowJo™ v10.8 Software (BD Life Sciences).

## Discussion

Antimalarial drugs that target apicoplast processes are widely used as prophylactics (e.g., doxycycline) and as components in second line combination treatments (e.g., clindamycin-quinine). The molecular target of most apicoplast inhibitors is the bacterial translation apparatus of the organelle. However, the means by which this leads to parasite killing through the delayed death phenomenon has not been thoroughly described. Interrogating downstream pathways of apicoplast metabolites, such as IPP, may reveal as yet unexplored drug targets. Direct and indirect inhibition of IPP production with either fosmidomycin or general apicoplast inhibitors is lethal, probably through the mis-prenylation of Rab proteins and other trafficking proteins. However, the fact that restoration of protein prenylation with GGOH does not lead to long term rescue of parasite viability (in contrast to IPP rescue) indicates that other isoprenoids are also essential for asexual RBC stage development. In this study we provide evidence that loss of apicoplast function in *P. falciparum* leads to a profound defect in dolichol synthesis and subsequently untethering of GPI-APs (**Fig 7**). Mislocalisation of GPI-APs leads in turn to 1) defects in merozoite segmentation during schizogony; 2) defects in merozoite ability to rupture the PVM and RBC membranes and failure to egress, and 3) defects in the ability of the small number of new merozoites that do form to invade new RBCs. Depletion of dolichol/polyisoprenoids lipids may also lead to disruption of parasite membrane structure and loss of viability.

**Fig 7.**
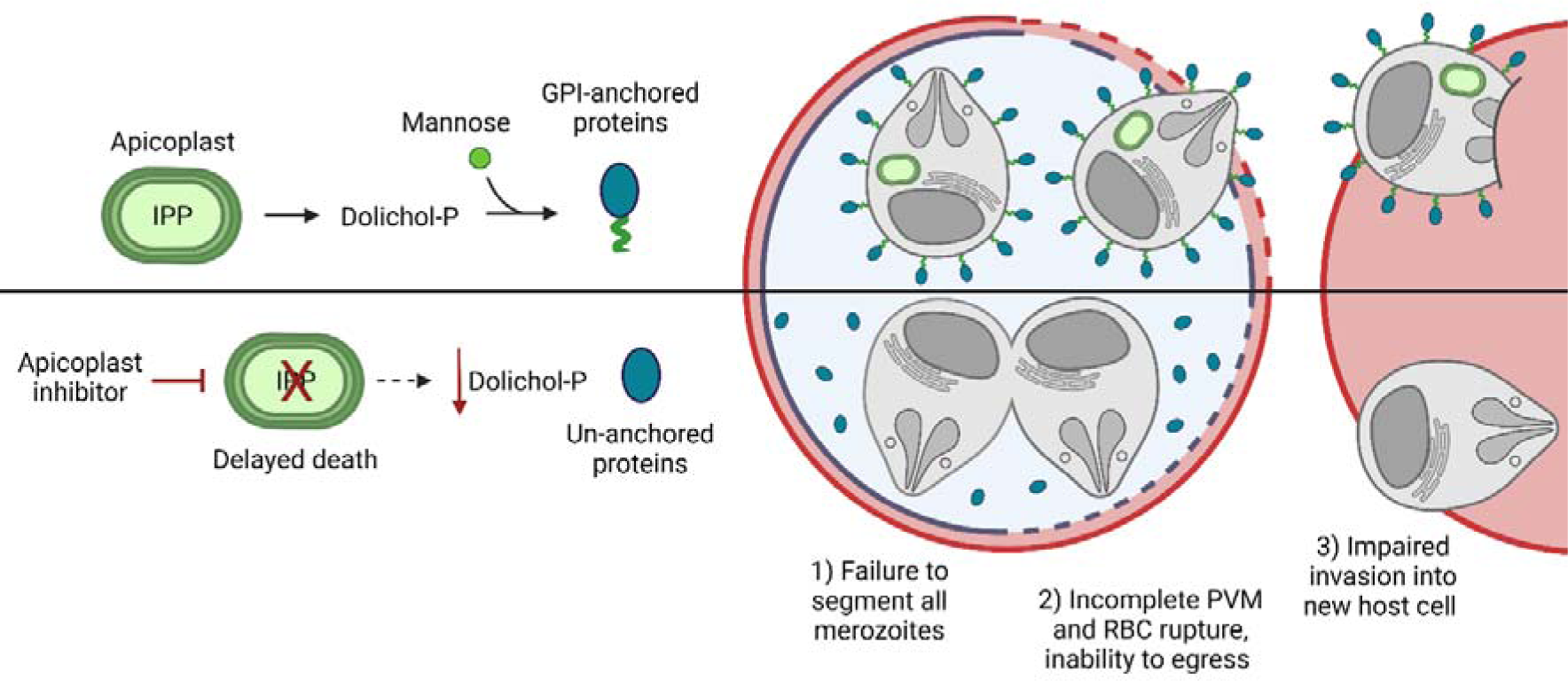
Dolichols are essential for survival of asexual stage *P. falciparum* parasites. Parasites treated with apicoplast inhibitors cannot synthesise IPP in their subsequent cycle. Without this essential metabolite, these parasites become depleted in dolichols which are required for the synthesis of GPI anchors and the normal membrane association of their anchored proteins. Upon mislocalisation of GPI-anchored proteins normal schizont development is halted with incomplete segmentation. These schizonts proved incapable of egress, and the merozoites that do form are unable to invade new RBCs even when mechanically purified from mature segmented schizonts.

During slow-onset depletion of IPP, as seen in delayed death, GPI-APs lose their membrane attachment in the second cycle, likely due to a block in dolichol synthesis and the depletion of the Dol-P-Man donor (**Fig 2A**). While a minority of GPI-APs treated with apicoplast inhibitors maintain their membrane anchorage, the gross mislocalisation of GPI-APs culminated in a lethal phenotype. Conversely, rapid inhibition of IPP synthesis by fosmidomycin left GPI-APs initially unaffected. This persistence of GPI-APs suggests that a low level of dolichols is sufficient for adequate GPI synthesis, as has been observed in other studies (23). *Plasmodium* parasites likely contribute to the maintenance of a surface glycocalyx on the merozoite plasma membrane, as well as acting as a large reservoir of GPI precursors (52). While these findings suggest that dolichol synthesis is essential for normal asexual RBC development, the apparent slow turnover of these lipids detracts from dolichol biosynthesis enzymes being good drug targets. In contrast, there is evidence that inhibition of enzymes involved in DPM or GPI biosynthesis can lead to rapid, first-cycle anti-malarial activity (53) (43, 54).

In addition to their roles in GPI biosynthesis and protein C-mannosylation, neutral polyisoprenol/dolichol lipids can also contribute to the biophysical properties of cellular membranes (55, 56). Intriguingly, *P. falciparum* asexual parasite stages produce unusually high levels of *cis*-polyisoprenols, which are present at the same concentration as mature dolichols (23). In most eukaryotes, levels of cis-polyisoprenoids are very low due to their conversion to dolichol by a polyprenol reductase. This enzyme is active in *P. falciparum* asexual stages but may be insufficient to convert the large steady state pool of cis-polyisoprenoid lipids to dolichols (23). Unlike dolichol, cis-polyisoprenols are not utilised as sugar carriers in eukaryotic cells and may therefore be primarily required for membrane function. In this respect, it is notable that parasites with GGOH-rescued apicoplasts exhibit abnormal IMC expansion and plasma membrane segmentation during merozoite segregation (**Fig 5F**). The factors and proteins that govern membrane invagination remain largely unknown, though sporozoite budding in the mosquito is at least in part regulated by the presence of the GPI anchor of circumsporozoite protein, possibly through the organisation of lipid-rich microdomains (57). GPI-APs in parasite membranes may play a role in these processes but free polyisoprenoids may also influence these membranes (23, 55, 56).

In addition to a segmentation failure, these PRDD schizonts were also incapable of egress (**Fig 5C**). This failure could be due to a block in a number of steps – our data showing only partial leakage of PV contents suggests at least one of these is a defect in PVM rupture. For parasites to become released into the bloodstream they must first break through two membranes; the PV and the RBC in a process which takes about 10 minutes in total (58). The current egress model suggests the PVM is initially permeabilised with localised pores that allow minor leakage of the PV contents into the RBC cytosol (59, 60). Shortly thereafter cGMP-dependent protein kinase G (PKG) is activated, which in turn triggers the secretion of the SUB1 protease into the PV (61). SUB1 cleaves a range of substrates within the PV and at the parasite surface, including MSP1. Next is the comprehensive disintegration of the PVM, the stage where we observe parasites stalling in PRDD treatment (**Fig 5D**). The specific effectors responsible for PVM breakdown are unknown, though this step can be blocked with C1, a PKG inhibitor. This suggests a SUB1 substrate may be responsible for PVM rupture, however MSP1 is the only known GPI-anchored substrate of SUB1 and appears uninvolved (41). For a GPI-AP to be involved in PVM rupture, it might regulate PKG or be activated by a SUB1 substrate. Alternatively, PVM disintegration may be multi-factorial, with a SUB1-independent pathway involving a GPI-AP also occurring. Once the PVM is fully ruptured, SUB1-activated MSP1 aids in the rapid destabilisation of the RBC cytoskeleton by binding to host spectrin. The mislocalisation of MSP1 from the merozoite surface in PRDD parasites would prevent egress at this step as well (**Fig 3B**). Our data implicate a yet uncharacterised GPI-AP with an essential role in egress during PVM rupture or perhaps a direct role of dolichols in PVM structure. Another possibility is that PVM rupture is mediated not only by parasite proteins, but by parasite-derived lipids as well. Neutral dolichols have been found in the lysosomal membrane of animal cells and thought to be essential for lysosome membrane integrity - it is an open question what eventually leads to a drain in dolichols following apicoplast inhibition and one possibility is that dolichols are transported to the PPM and subsequently to the PVM (e.g., via NPC1).

Despite incomplete segmentation in PRDD parasites, some merozoites were still formed in these conditions. However, these merozoites were also defective, being unable to invade new RBCs presumably due to a lack of GPI-APs at the parasite surface (**Fig 6D**). During normal invasion, the merozoite is completely coated in GPI-APs which act as a sticky interface with the RBC. These initial interactions are weak and reversible, allowing the merozoite to re-orient itself so the apical pole is facing at the RBC surface. The apical organelles then discharge proteins which mediate higher affinity binding and the formation of the tight junction, which becomes the entry point through which the merozoite actively traverses into the RBC. There are several GPI-APs at the merozoite surface, and it is unclear whether they bind distinct ligands or if there is functionally redundancy. In addition to its role in egress, MSP1 can form a variety of complexes with other MSPs (MSP3, MSP6, MSP7, MSPDBL1, MSPDBL2). Peptides from both MSP1 and its partners have shown RBC binding at glycophorin A and band 3, and inhibiting these interactions results in reduced merozoite invasion (62-67). While other GPI-APs such as MSP4, MSP8, MSP10, Pf12, Pf34, Pf38, Pf92, GPI-anchored micronemal antigen (GAMA) and surface related antigen (SRA) are less well characterised, incubating parasite cultures with their peptides or antibodies against these proteins also reduces invasion. Therefore, the invasion failure observed in our merozoites likely reflects the cumulative loss of GPI-APs and reinforces GPI biosynthesis as a potential drug target. While chemically targeting one GPI-AP may run the risk of functional compensation by other GPI-APs, inhibiting overall GPI-synthesis negates this possibility. Of the apical GPI-anchored proteins, knock-down of the GPI-anchored protein rhoptry associated membrane antigen (RAMA) results in defective rhoptry neck formation and consequently invasion, presumably due to impaired protein discharge (68). While the localisation of RAMA was not addressed in our study, we found no apparent morphological defect in rhoptry neck and bulbs. Whether this reflects maintained RAMA anchorage and/or trafficking is unclear. However, we predict these merozoites are incapable of re-orientation and therefore rhoptry discharge still may not occur.

We have shown that apicoplast derived IPP is essential for the normal anchorage and localisation of GPI-anchored proteins in blood stage *P. falciparum*. The dolichol required for GPI synthesis are likely depleted concurrently with or after the exhaustion of apicoplast synthesised IPP. While defects in protein prenylation is the proximate cause of death of parasites treated with delayed-death drugs, we show that additional consequences of such drugs include a block in GPI synthesis, impacting parasite segregation, egress, and invasion. This study has revealed novel roles for GPI-APs in schizont segmentation and rupture of the PVM, as well as confirming the importance of these proteins for RBC rupture and merozoite invasion. The wide-reaching consequences in parasites with depleted dolichols strongly support GPI-synthesis as a novel anti-malarial target.

## Acknowledgements

We are grateful for the generous donations from Matthew Dixon for the AMA1 antibodies, Robin Anders for the various MSP2 antibodies (39), Alan F. Cowman for the GAP45 antibodies and Paul Gilson for the MSP1 and EXP2 antibodies, along with the PEXEL-Nluc cell line. For invaluable technical advice we thank Paul Gilson and Madeline Dans in regard to the egress assays. Images created with Biorender.com. MJM is a NHMRC Principal Research Fellow. This research was supported by a grant from the NHMRC (1181336)

