## Supplemental information for "Apicoplast-derived isoprenoids are essential for biosynthesis of GPI protein anchors, and consequently for egress and invasion in *Plasmodium falciparum*"

Supporting Information:

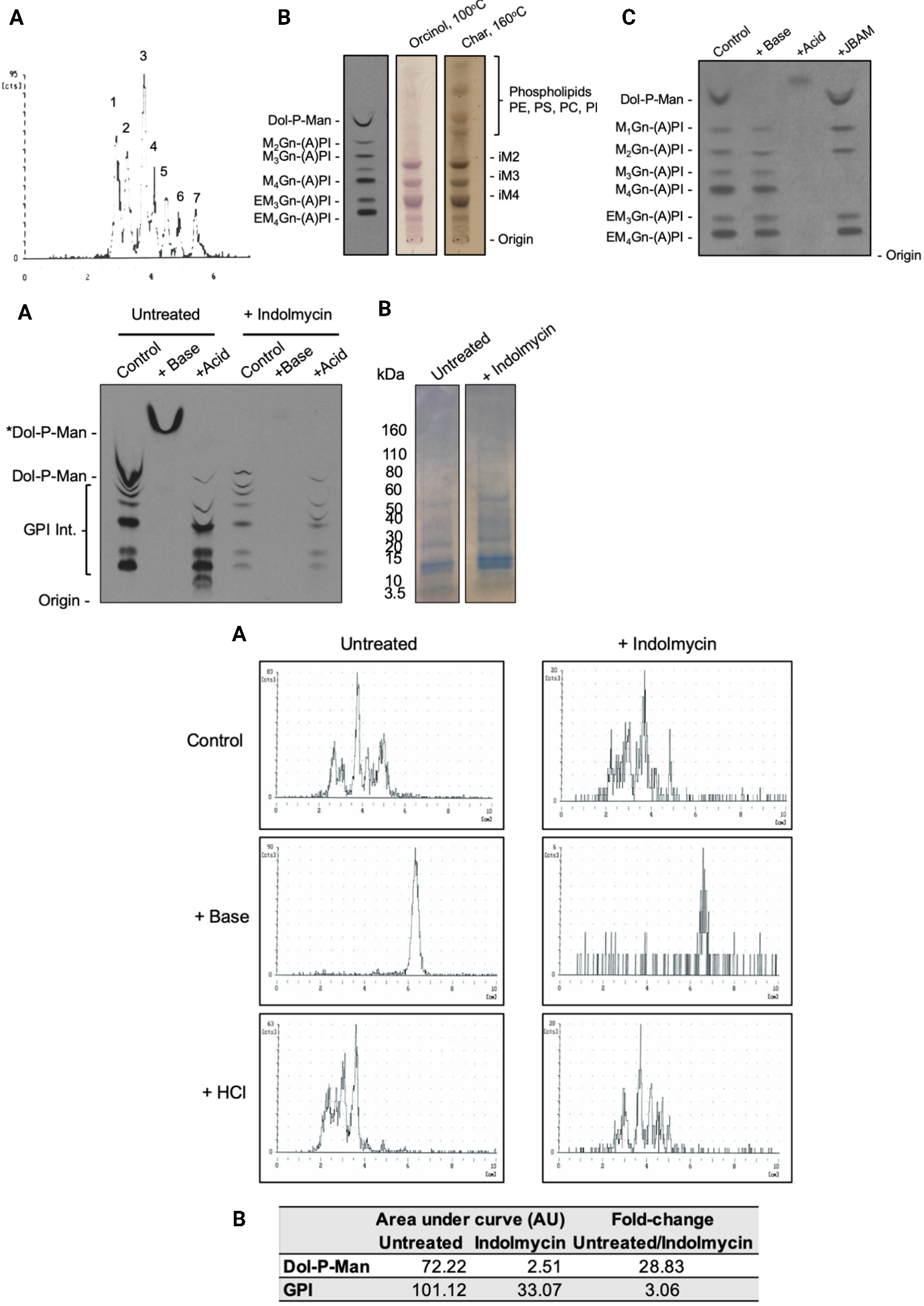

**S1 Fig.**

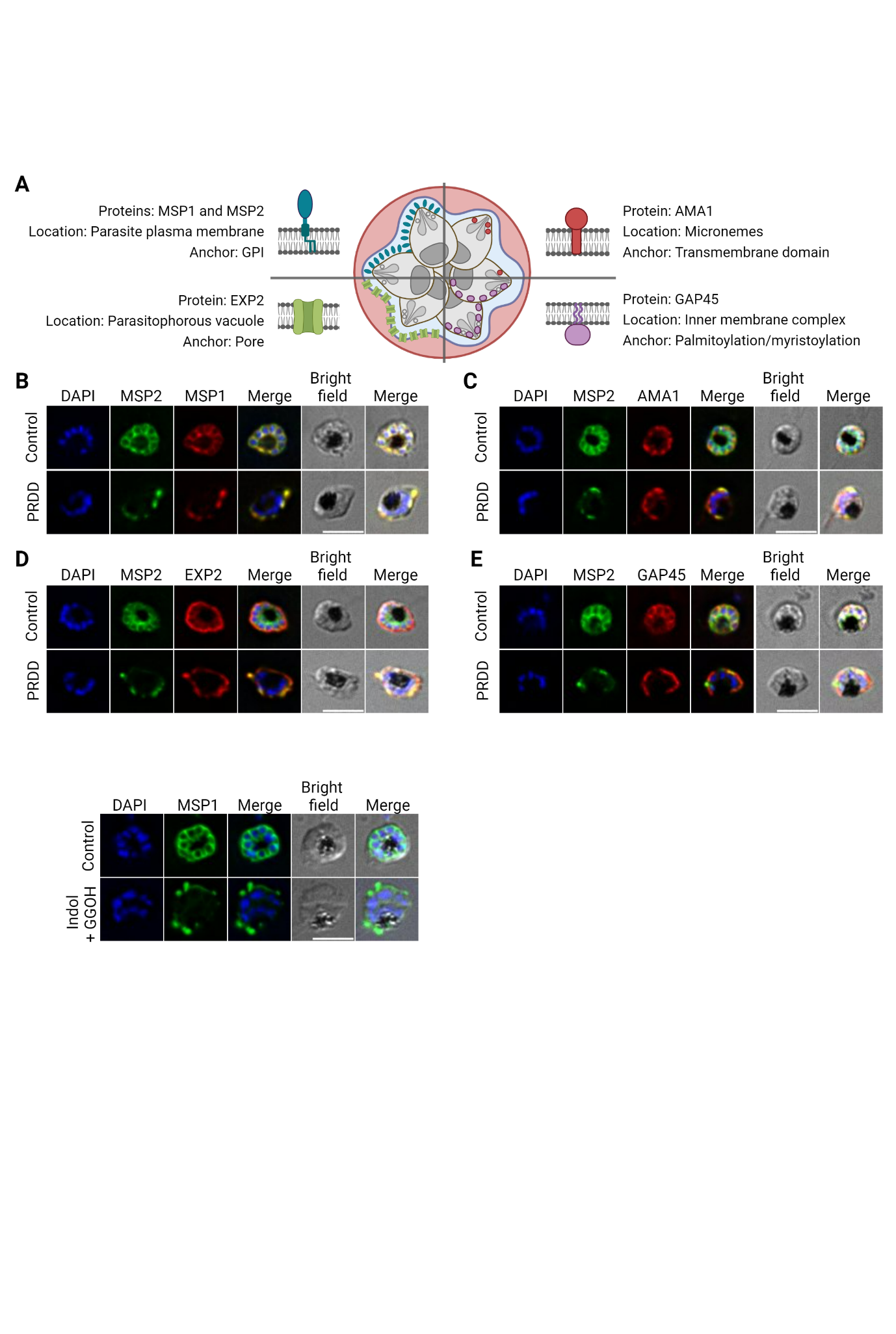

**S2 Fig.**

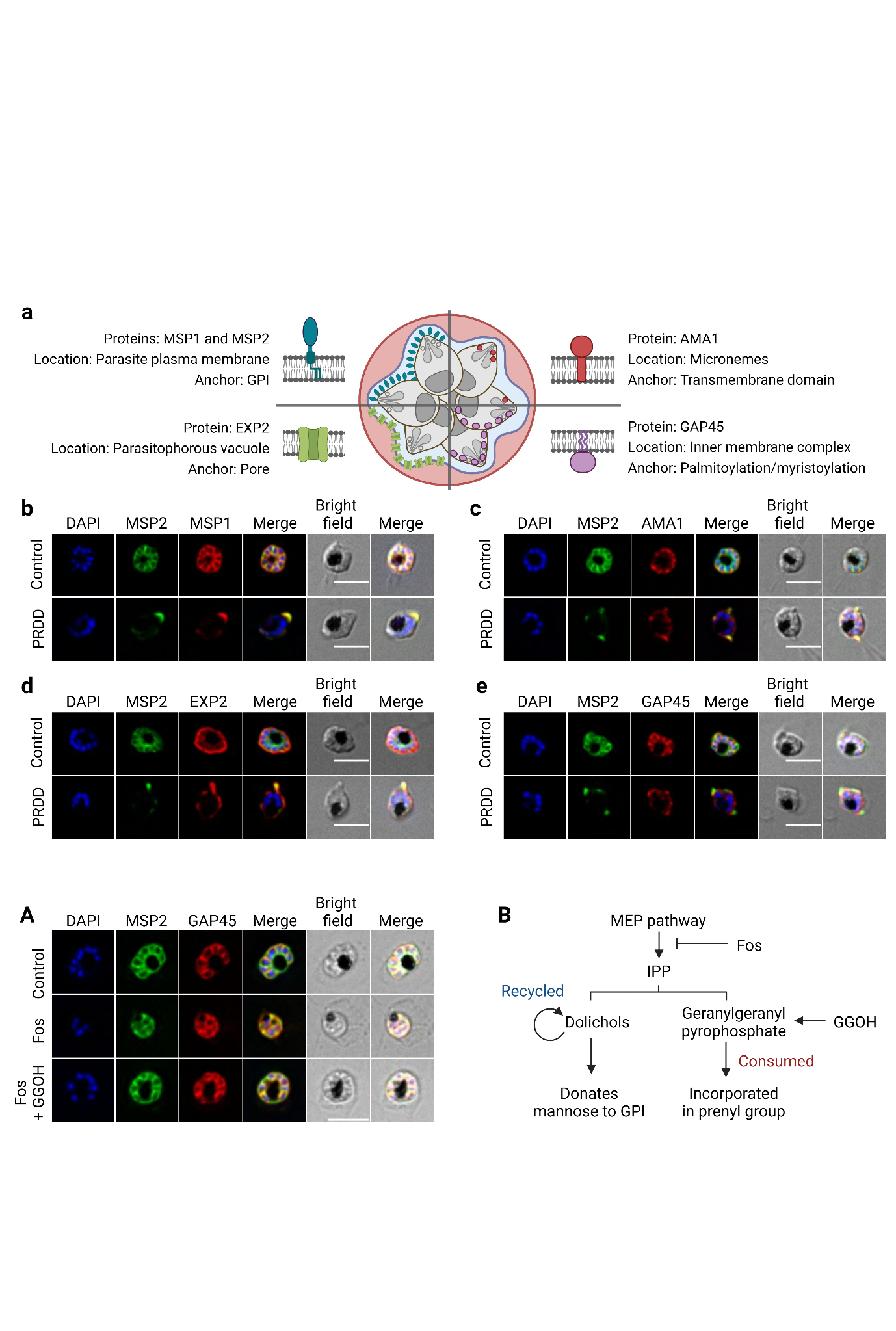

**S3 Fig.**

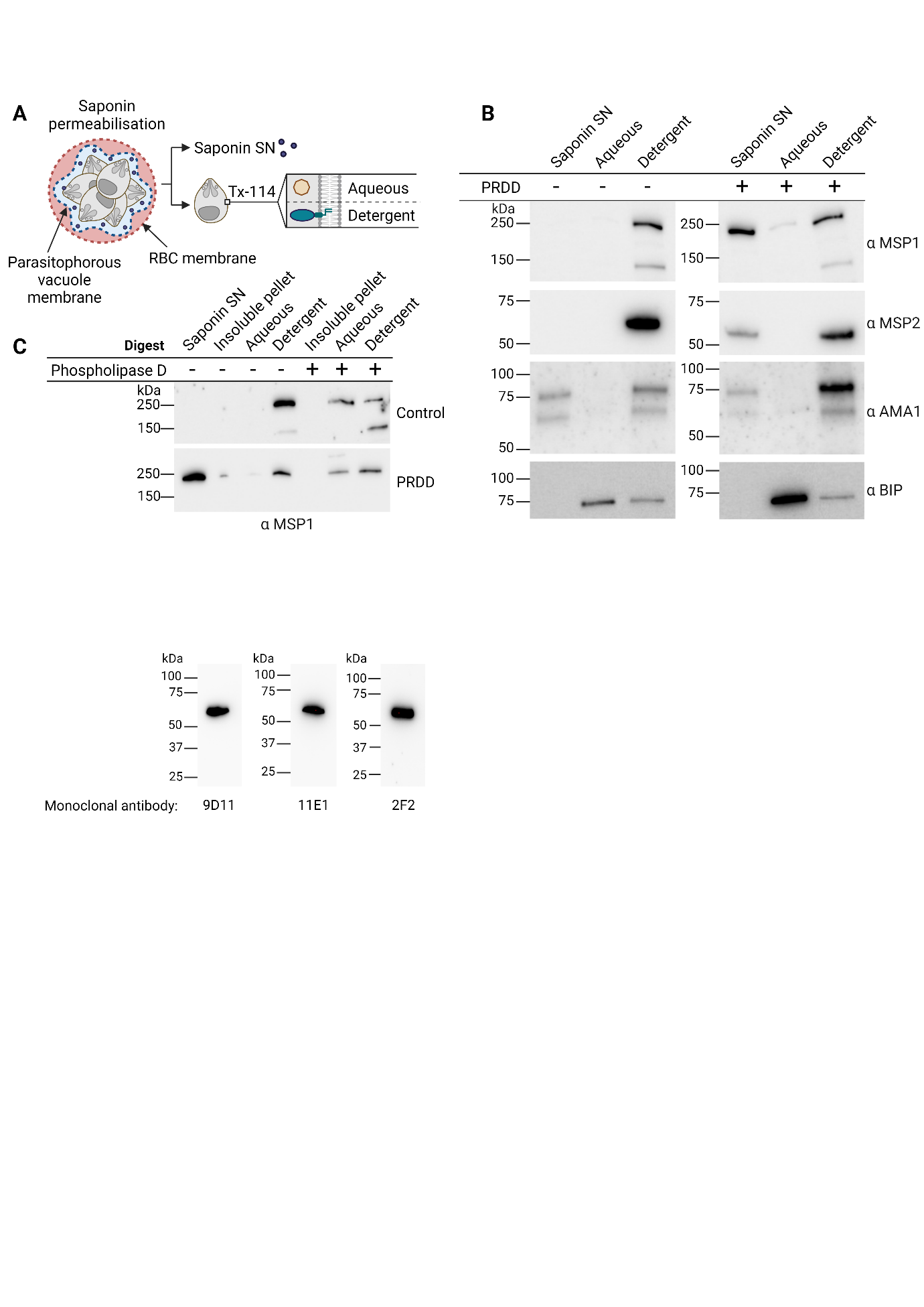

**S4 Fig.**

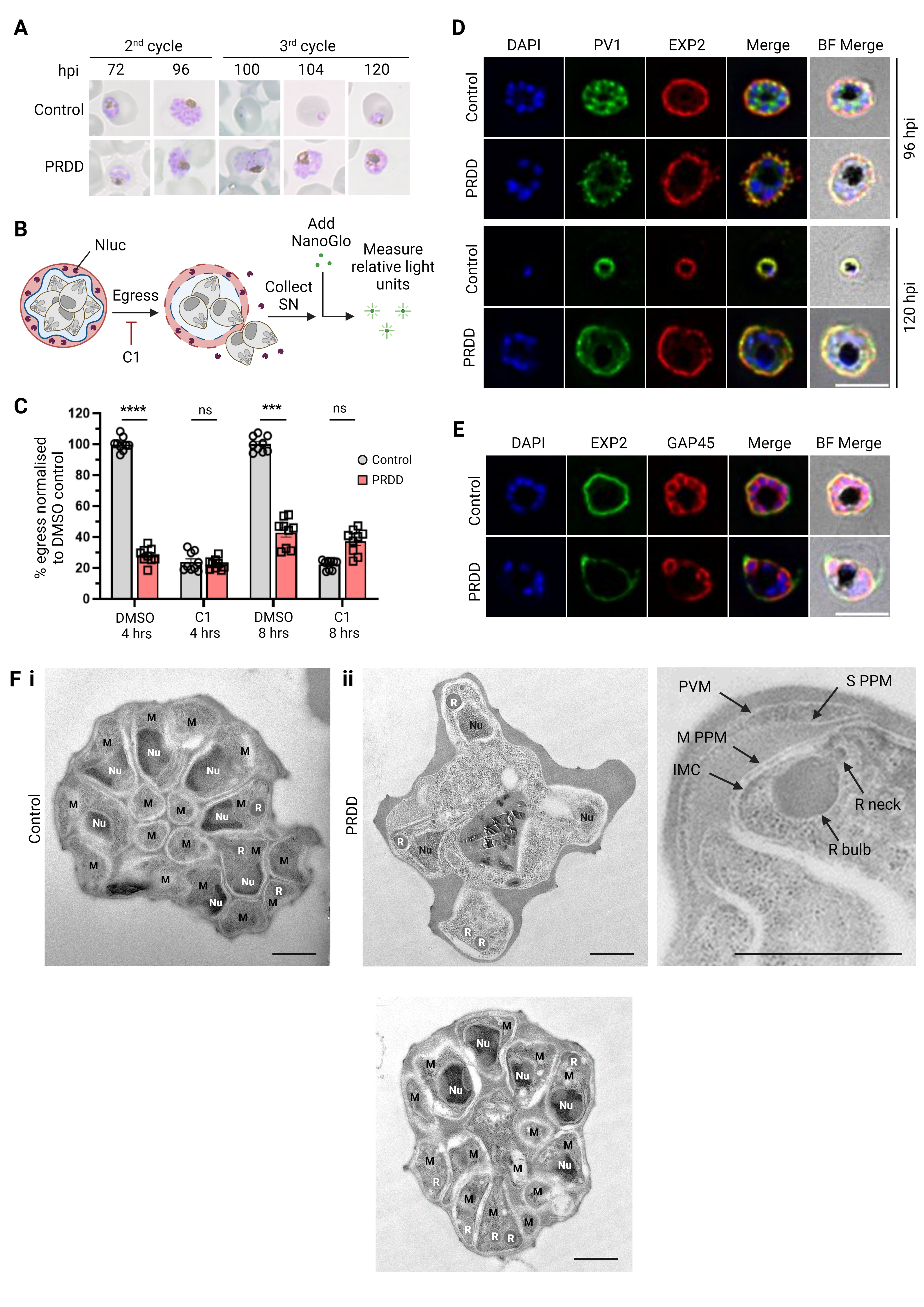

**S5 Fig.**

**S1 Fig. Linear scan of HPTLC containing [3H]-man incorporated P. falciparum glycolipids with chemical treatments. A)** Distance (cm) is shown on the x-axis and counts per minute (CPM) on the y-axis. **B)** The total area under the curve (equivalent to proportion of [^3^H]-man incorporation into) was estimated to determine the fold-change differences between abundance of glycolipids in untreated and treated conditions. AU, arbitrary units.

**S2 Fig. PRDD induced with indolmycin also results in mislocalisation of surface GPI-anchored proteins.** Treatment with apicoplast translation inhibitor indolmycin (Indol) with GGOH results in the mislocalisation of surface GPI-anchored protein merozoite surface protein 1 (MSP1). Nuclei stained with 4,6-diamidino-2-phenylindole (DAPI; blue). Scale bar = 5 µm.

**S3 Fig. Fosmidomycin treatment rapidly inhibits synthesis of prenyl groups but does not immediately impact GPI-anchoring of proteins. A)** Fosmidomycin (Fos) treatment is lethal, killing at the trophozoite stage. Addition of geranylgeraniol (GGOH) allows parasites to progress to the schizont stage where GPI-anchored MSP2 (green) maintains surface localisation. Nuclei stained with DAPI (blue). Images represent single Z stacks. Scale bar = 5 µm. **B)** Dolichol and geranylgeranyl pyrophosphate are both synthesised from apicoplast-derived IPP. Dolichols are recycled in their role as sugar donor and so are more robust to IPP inhibition. Geranylgeranyl pyrophosphate is consumed to form the prenyl group itself and following fosmidomycin treatment is quickly depleted with lethal consequences.

**S4 Fig. Merozoite surface protein 2 (MSP2)** **migrates at a higher apparent molecular weight than predicted.** Western blot of 3D7 parasite lysates probed with anti-MSP2 monoclonal antibodies show MSP2 consistently migrating at a size larger than the predicted 28 kDa.

**S5 Fig. Some parasites with normal segmentation are also observed during delayed death.** Schizonts with clear and total segmentation of nuclei (Nu), rhoptry (R) formation and membrane encapsulated merozoites (M) can be seen, despite drug treatment. Scale bar = 1 µm.
